# High *Plasmodium* infection intensity in naturally infected malaria vectors in Africa

**DOI:** 10.1101/780064

**Authors:** Anais Bompard, Dari F. Da, Serge R. Yerbanga, Isabelle Morlais, Parfait H. Awono-Ambéné, Roch K. Dabiré, Jean Bosco Ouédraogo, Thierry Lefèvre, Thomas S. Churcher, Anna Cohuet

## Abstract

The population dynamics of human-to-mosquito malaria transmission in the field has important implications for the genetics, epidemiology and control of malaria. The number of oocysts in oocysts positive mosquitoes developing from a single, naturally acquired infectious blood meal (herein referred to as parasite exposure) greatly influence the effectiveness of transmission blocking interventions but still remains poorly documented. During a year-long analysis of malaria parasite transmission in Burkina Faso we caught and dissected wild malaria vectors to assess *Plasmodium* oocysts prevalence and load (the number of oocysts counted in mosquitoes with detectable oocysts) and the prevalence of salivary gland sporozoites. This was compared to malaria endemicity in the human population assessed in cross-sectional surveys. Data was analyzed using a novel transmission mathematical model to estimate the per-bite transmission probability and the average parasite exposure of mosquitoes for each location. Observed oocysts load and estimated parasite exposure in naturally infected mosquitoes is substantially higher than previous estimates (ranging from 3.2 to 24.5 according to seasons and locations) and indicates a strong positive association between parasite exposure of mosquitoes and parasite prevalence in human. This work suggests that highly infected mosquitoes may have a greater influence on the epidemiology and genetics of the parasite and that novel partially effective transmission blocking interventions may become more effective at halting transmission as parasite exposure is diminished.

## Introduction

In the effort to control malaria, understanding the processes governing transmission of wild parasites between vectors and vertebrate hosts is paramount. Quantitative information on the number of parasites in wild mosquito populations is lacking and very few studies have investigated either the infectivity of the human parasite reservoir to wild mosquitoes (1–6) or malaria parasites in naturally infected wild mosquitos(7–11).

Oocyst intensity describes the average number of visible oocysts in all mosquitoes fed on the same blood-source(12). This quantity is easy to measure in experimental infections (where lab-reared mosquitoes can be given a single blood meal and maintained in the insectary) though it is difficult to quantify in the wild. This is because the number of detectable oocysts in wild caught mosquitoes may have been acquired over multiple blood meals and be at various stages of development. For example, wild mosquitoes might feed multiple times before oocysts developing from gametocytes ingested during the first blood meal become observable. Here we define the term oocyst load to be the average number of oocysts in wild caught mosquitoes with detectable oocysts (by microscopy), and parasite exposure to be the average number of oocysts acquired during a single infectious blood meal that leads to oocyst development. Oocyst intensity, load and exposure are related to one another but only oocyst load is directly measurable in wild caught mosquitoes as the sources of previous blood meals are unknown.

Understanding the effectiveness of current and novel drugs and vaccines which targets malaria and how best they should be evaluated and deployed in the field depends on a better understanding of human populations infectivity(13–15). Vaccines which interrupt malaria parasite transmission by targeting sexual and sporogonic stages of the parasite are referred to as transmission-blocking vaccines. The efficacy of these vaccine candidates in reducing the number of infected mosquitoes shows a dose response to parasite exposure(16–18) which depends primarily on the number of mature gametocytes in the blood, though many other factors are known to contribute. These include gametocyte maturity, parasite clone diversity, host blood component and environmental conditions (18–23). Parasite exposure therefore represents the realised infectiousness of a human host assuming that the mosquito survives long enough for all oocysts to develop.

The current consensus is that oocyst load and parasite exposure (the number of oocysts developing from a naturally acquired infectious blood meal) in the wild is very low, with each wild mosquito carrying less than 5 oocysts(24). This may be true but very few studies have directly investigated parasite exposure and values are likely to vary between locations and over seasons as environmental conditions and host factors affect the endemicity of *Plasmodium* transmission (7,25). Field studies involving collection and dissection of wild caught mosquitoes are logistically difficult and time-consuming so it is not practical to repeat these studies every location where a transmission blocking interventions may be used. Thus, to predict the intervention efficacy in the field and plan for deployment, we need to better understand the level of parasite exposure in the wild and find easier-to-collect measures of disease endemicity capable to predict its changes.

A year-long field study was conducted in two high transmission villages in Burkina Faso, collecting information on *Plasmodium* parasites in wild mosquitoes caught in huts and in the local human population. Wild mosquitoes were also collected and dissected to investigate for *Plasmodium* parasites in two villages of Cameroon. It is not possible to reliably quantify the number of oocysts in a mosquito by dissection until the latest blood meal has digested, therefore mosquitoes collected in huts were kept in the insectary and dissected 3 and 7 days following collection(8) to detect oocysts and sporozoites. A novel mosquito transmission model was developed to support data analysis and estimate parasite exposure from the collected data. Results indicate that oocyst load and parasite exposure in high transmission areas were much higher than usually assumed and can be broadly explained by gametocyte prevalence (and to a lesser extent by all stages parasite prevalence) in the human population.

## Results

### 1. Parasites in humans

Prevalence for *Plasmodium falciparum* as determined by microscopy was particularly high in the study villages in Burkina Faso, with between 22-32% infected people during the dry season (collections in March and April) and 35-55% during the rainy season (collections from July to November), depending on month and location (figure 1. a, b). Asexual parasite prevalence was estimated from ~300 individuals at each time point and was significantly higher in Klesso than in Longo (dry season 32.3% (95% confidence intervals: 27-38%) vs. 22% (17-26%), p-value= 0.003; rainy season: 49% (45-52%) *vs*. 44% (40-47%), p-value= 0.042), though observed gametocyte prevalence were similar between locations. Both asexual parasite and gametocyte prevalence varied across seasons, reaching a maximum in September (at the peak of the rainy season) and a minimum during the dry season for asexual parasites and in November for gametocytes. The density of asexual parasites varied across season and location (figure 1. a, b, supporting table 1), but gametocyte density was independent of seasonal variations and similar between localities (supporting table 1).

**Figure 1.**
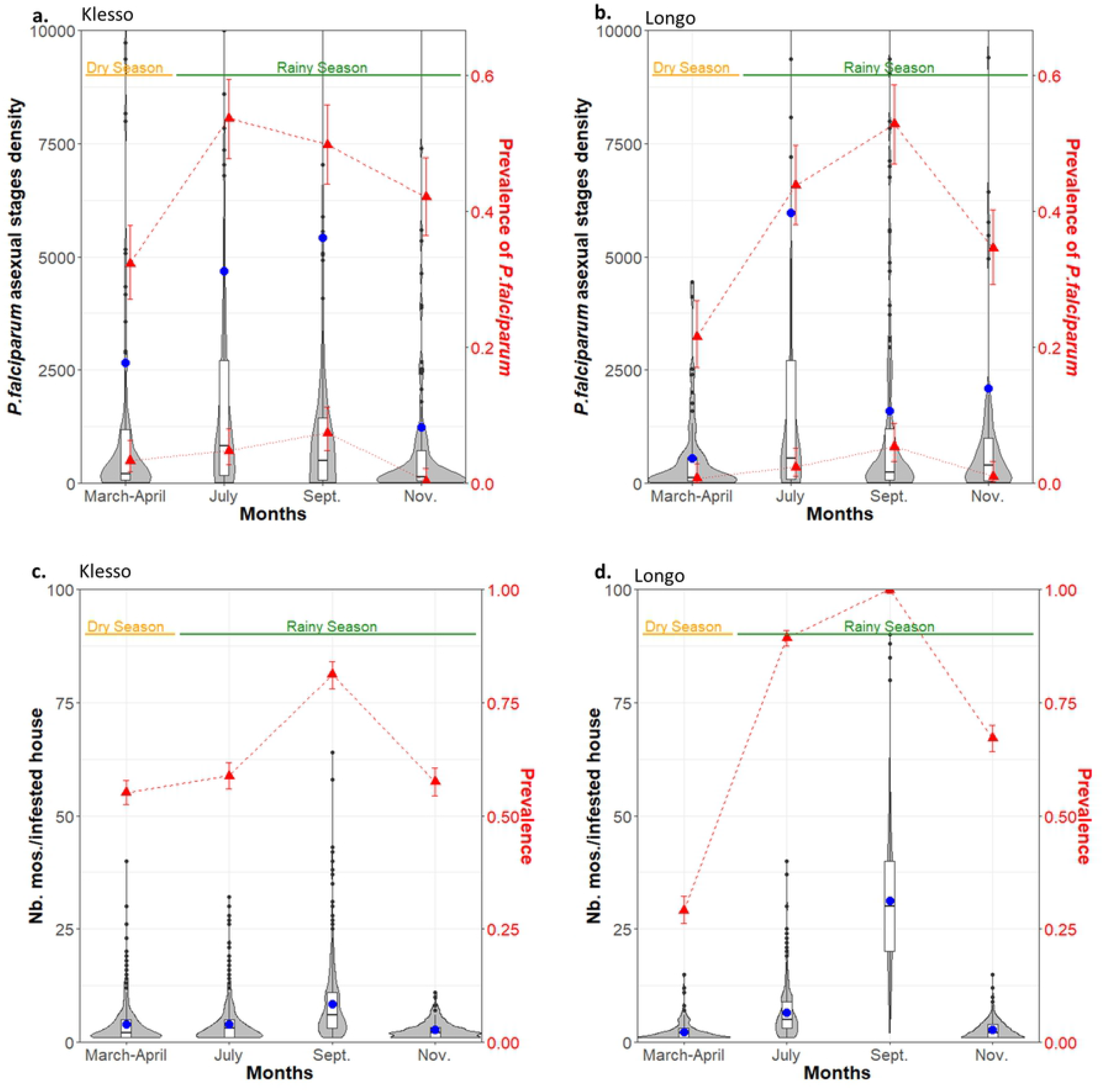
Human parasite prevalence and density (top row) and mosquito abundance (bottom row) from two villages in Burkina Faso (Klesso = first column, Longo = second column) collected at 4 time points throughout the year. a-b: *Plasmodium falciparum* asexual parasite density (trophozoite, measured as parasites/μl) is shown on the left axis. Violin plot and point and whisker plots show the population density distribution with the mean as a blue dot. Right axes and red points show parasite prevalence, be it all stage parasites (triangular points, dashed line) or gametocytes (triangular points, dotted line) as measured by microscopy. c, d: Number of mosquitoes (all genus) per infested houses (left axis: violin plot with mean as a blue dot) and prevalence of mosquitoes in houses (right axis: red line and points).

Prevalence of *P falciparum* infections also varied with age class (0-5, 5-20, 20-50, and >50 years-old), with highest asexual and gametocyte prevalence in the 0-5 years-old (61% (52-70%) asexual parasite carriers and 7% (2-11%) gametocyte carriers) and lowest for the 20-50 years-old (see supporting table 1, supporting figure 1). *P. falciparum* composed over 98.9% (98.5-99.3%) of all *Plasmodium* infections, the remaining infections being split between *Plasmodium ovale* and *Plasmodium malariae*.

### 2. Mosquito abundance

The prevalence and abundance of mosquitoes in households, including all collected genus, varied substantially throughout the year and between locations in Burkina Faso (Figure 1. c, d), as did the mean number of *Anopheles* mosquitoes per household (supporting table 2, supporting figure 2).

Species composition among collected *Anopheles* differed between locations: *Anopheles gambiae s.s*. was the most prominent species in Klesso (55.8% (52-60%)) whereas *Anopheles coluzzii* was the most common in Longo (62.9% (59-67%), see supporting table 2). *A. coluzzii* was the prominent *Anopheles* mosquito collected during the dry season at all locations (80% (75-85%)). Gravidity, parturity rates and feeding status of mosquitoes were also reported (supporting table 2). At the start of the rainy season, we found significantly more parous mosquitoes in Klesso than in Longo, suggesting a larger proportion of older mosquitoes in accordance with their observed presence during the dry season.

Bednet coverage was significantly higher in Longo than in Klesso all year round (85.1% (84.0-86.2%) vs. 56.9% (55.3-58.4%), p-value<0.0001, see supporting table 2, supporting figure 3). Bednets were likely used in response to mosquito nuisance, as bednet coverage followed mosquito prevalence and abundance over time in each location. During the peak of the rainy season, bednets significantly reduced the number of mosquitoes in infested houses at all locations (−15.8%, p-value<0.0001, supporting figure 3). Other houses characteristics (type of roof, presence of paint) were reported but had a very limited impact on mosquito prevalence and abundance (supporting table 2).

### 3. Parasites in mosquitoes

Wild mosquitoes caught on the inside of houses were kept for three days (D3) or seven days (D7) in the insectarium in order to allow sufficient time for the last blood meal to be digested, enabling oocyst load to be quantified by microscopy after dissection. The difference in oocyst prevalence and abundance between D3 and D7 allows an evaluation of the transmission dynamics (see materials and methods for more details).

#### Oocyst prevalence and oocyst load at D7

The prevalence of oocyst-infected mosquitoes at 7 days after collection varied significantly with seasons (p-value<0.0001), with a maximum in September (25.9% (23.0-28.8%) in Klesso and 8.1% (6.4-9.9%) in July in Longo). During the dry season prevalence dropped to 5.0% (2.5-7.2%) in Klesso and no oocysts were detected in Longo (figure 2. a, b). Average oocyst load among infected mosquitoes (the number of detectable oocysts in the midgut) during the rainy season was similar in both villages at D7 with 13.8 (10.1-17.1) oocysts per mosquito in Klesso and 10.3 (7.8-12.5) oocysts/mosquito in Longo (p-value=0.73). Oocyst load varied with season (p-value<0.0001), reaching 14.3 (7.0 – 19.7) oocysts/mosquito in September in Klesso and 14.7 (8.9 – 20.3) in Longo, figure 2. a, b). Oocyst distribution within mosquitoes was highly overdispersed (aggregated) with a median oocyst load of 3 oocysts/mosquito but 5.8% of infected mosquitoes presenting over 50 detectable oocysts (56 out of 977 mosquitoes examined). Two percent of infected mosquitoes had over 100 detectable oocysts (figure 2 c, d). Mosquito species was not significantly associated with oocyst prevalence (p-value = 0.23) or load (p-value= 0.26).

**Figure 2.**
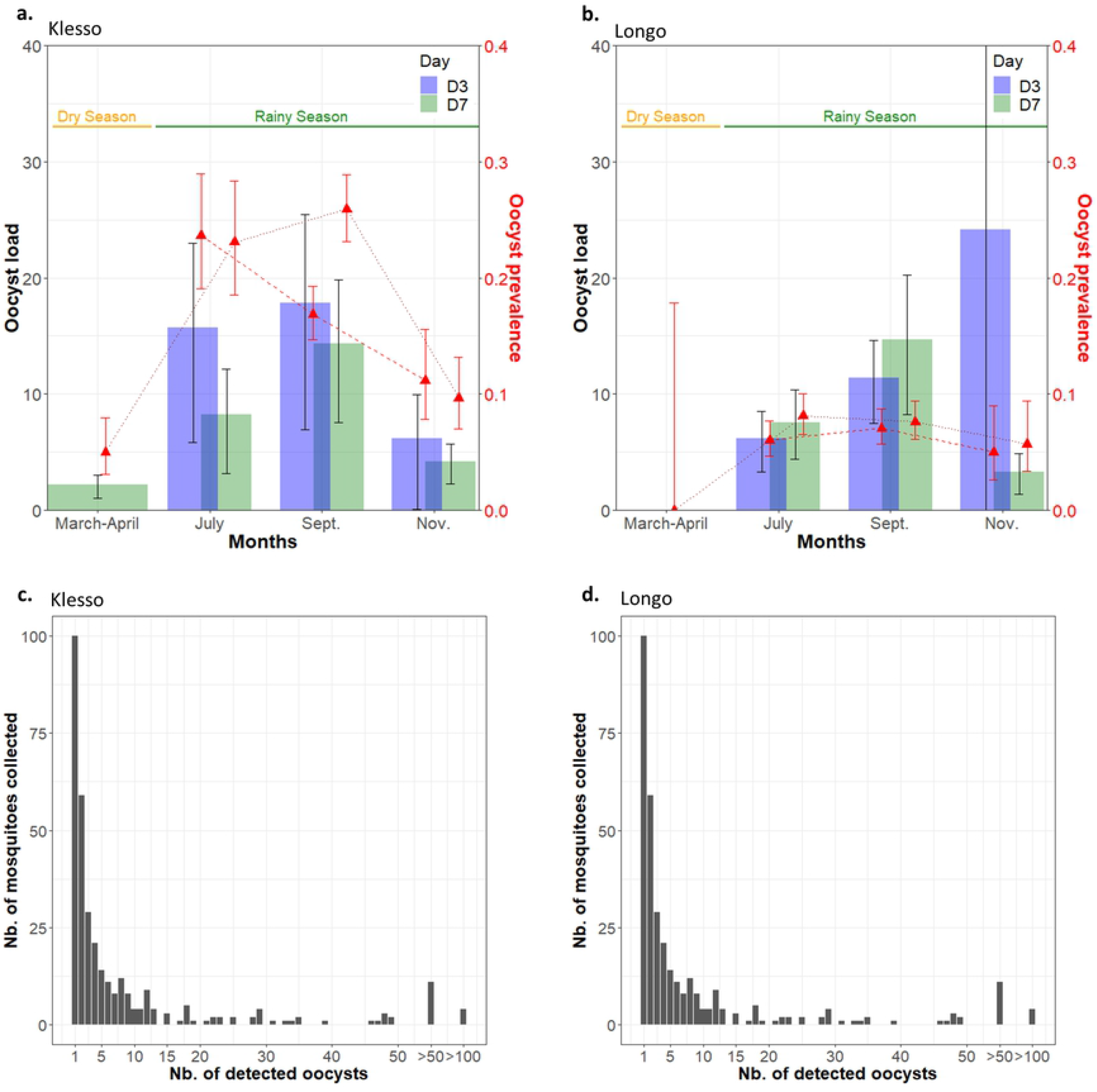
*P. falciparum* oocyst prevalence and load in mosquitoes from the villages of Klesso (first column) and Longo (second column). a, b: Mean oocyst load (average number of oocysts per mosquito with detectable oocysts) 3 days (blue) or 7 days (green) after mosquito collection (barplot with 95% confidence intervals) and prevalence of oocysts 3 days (D3) and 7 days (D7) after infection (right axis: blue triangle and dashed red line (D3) or green triangle and dotted red line (D7). No dissections were done at D3 during the dry season). c, d: The distribution of number of oocysts per infected mosquitoes (bar for uninfected mosquitoes has been omitted) measured 3 or 7 days after blood-feeding.

Visible shells of oocysts that have already ruptured were also detected but this information was not included in the transmission model due to uncertainty in their persistence on the mosquito midgut after the release of sporozoites (see supporting information 1). During the rainy season 3.4% (2.2-4.5%) of mosquitoes presented ruptured oocysts, among them the average of ruptured oocysts was 13.2 (10.3 – 15.8) and one mosquito displayed 127 detected ruptured oocysts.

#### D3-D7 changes and D0 estimates

Oocyst prevalence 3 days after collection was not significantly lower than 7 days after collection (Longo: p-value= 0.098, Klesso: p-value= 0.86) as would have been expected if oocysts are undetectable 3 days after the infectious blood meal(8). Oocyst load even significantly decreased between D3 and D7 in Klesso (−24.9%, p-value= 0.013) while it displayed no significant difference in Longo (p-value= 0.61) (Figure 2. a, b). In order to understand the evolution of oocyst prevalence and load between D3 and D7, we monitored oocyst threshold of detectability in experimentally infected mosquitoes in the laboratory (see supporting information 2). This supporting experiment indicated that a substantial proportion of oocysts were already detectable 3 days after an infection: the prevalence of oocysts in mosquitoes dissected at day 3 was on average 60% of their prevalence at day 7 (with a range of 0 to 100% depending on the blood donor). The proportion of mosquitoes with detectable oocysts increased with parasite exposure (measured as the number of oocysts at day 7 after infection, as only one blood meal was taken and all oocysts are considered developed at this time). This feature was then included in the transmission model (see supporting information 2 and 3).

#### Sporozoite rates

Along the the rainy season, from July to November, the prevalence of infectious mosquitoes (with sporozoites) 7 days after collection ranged between 27-38% in Klesso and between 12-34% in Longo (significantly lower in Longo, p-value<0.0001, figure 3 a, b). In both locations, sporozoite prevalence reached its maximum at the end of the rainy season. Prevalence of infectious mosquitoes was significantly lower at D3 after collection (compared to D7, p-value<0.0001), ranging between 7-27% in Klesso and 11-23% in Longo across the season. We used the transmission model described in supporting information 3 to estimate the normalised prevalence of infectious mosquitoes in the wild, as would have been detected if the mosquitoes had been dissected on the day of collection: these estimates range between 2-17% in Klesso and 2-7% in Longo (figure 3 a, b). Mosquito species within the *Anopheles* group had no impact on sporozoite prevalence (p-value = 0.672).

**Figure 3.**
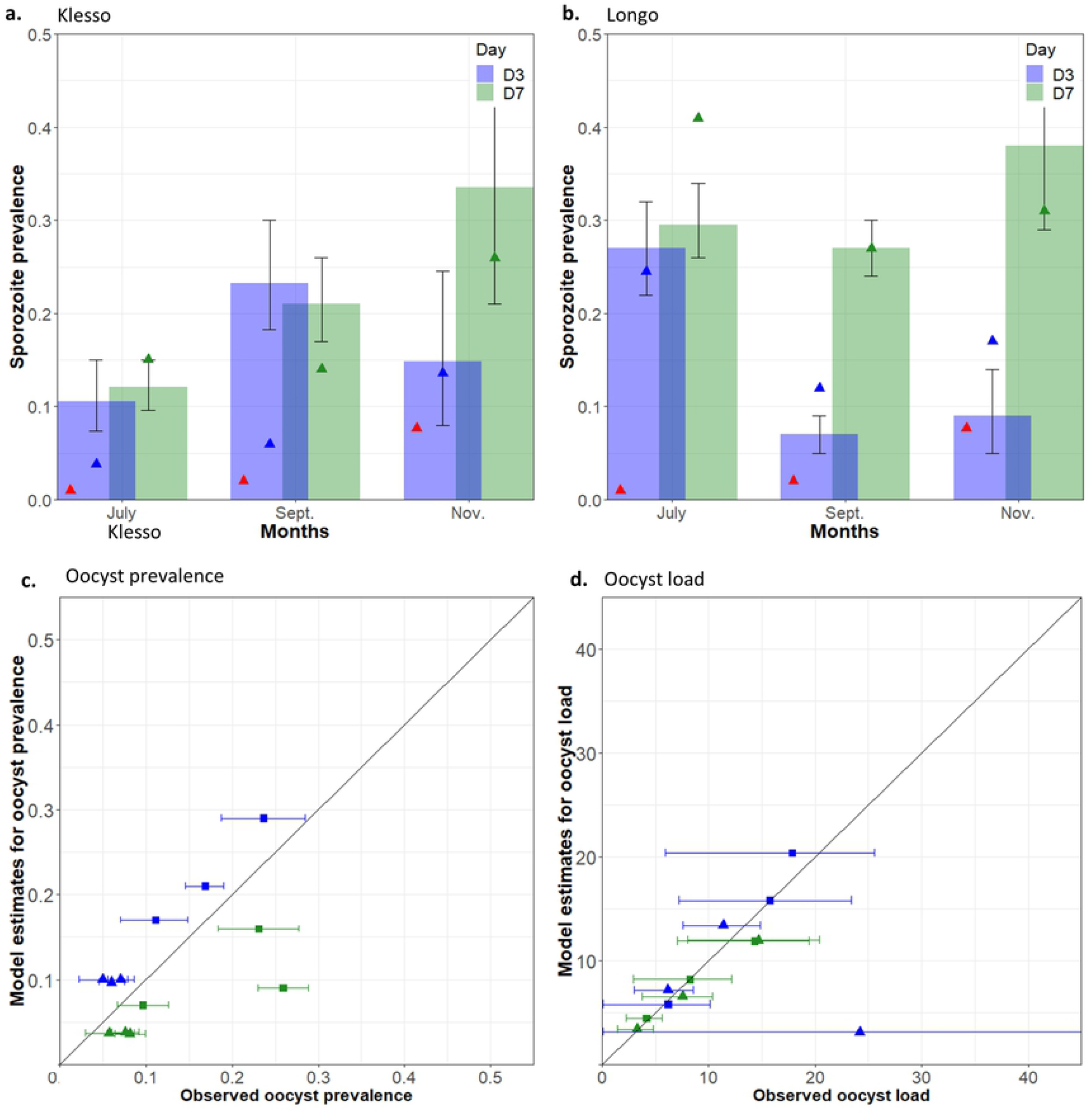
Observed data on sporozoite prevalence and model predictions for *P. falciparum* in mosquitoes in Kesso and Longo. a, b: Observed sporozoite prevalence at D3 (blue bars) and D7 (green bars) with vertical black lines indicating 95% confidence interval estimates. Model prediction for sporozoite prevalence in the mosquito population at the collection date (red triangles), at D3 (blue triangles) and D7 (green triangles) for Klesso (a) and Longo (b). c, d: The ability of the model to capture oocyst prevalence and load. c: Model prediction for oocyst prevalence at D3 (blue) and D7 (green) vs. observed oocyst prevalence at these times (with 95% confidence intervals on data) for Longo (triangles) and Klesso (squares). d: Model prediction for oocyst load (the mean number of oocysts per mosquito) at D3 (blue) and D7 (green) vs. observed oocyst load at these time (with 95% confidence intervals)) for Klesso (squares) and Longo (triangles).

#### Infection rate, parasite exposure and superinfections

The infection rate (the probability of transmission between human and mosquito per blood meal on an infected host) and parasite exposure (the number of oocysts developing in a mosquito following a single infectious blood meal) were estimated during the rainy season by fitting a Susceptible-Exposed-Infectious compartmental deterministic *Plasmodium* transmission model to data on human and mosquito parasite prevalence and load (3 and 7 days after mosquito collection). Details of the mathematical model are provided in supporting information 3 with results presented in figure 3 and supporting figure 4. Estimates of the probability of human-to-mosquito transmission (per blood meal taken on a person with detectable parasites by microscopy) showed little variation between seasons with an average estimate of 35% across all timepoints. Values were higher in Klesso (39 – 79%) than in Longo (9 – 15%) (see supporting information 3). Overall estimates of parasite exposure across all villages and timepoints showed that infectious blood meal resulted in an average of 14 oocysts per mosquito. Mean parasite exposure was consistent between villages, with estimated number of 7.0-24.5 oocysts/infectious blood meal in Klesso and 3.2 – 18.3 oocysts/infectious blood meal in Longo. Exposure appeared to vary by season, being highest at the peak of the rainy season (September) and a minimum at the end of the rainy season (November) in both locations. Mosquito average life-expectancy as estimated by the model was similar between locations (around 8 days in Klesso, 7 days in Longo), and we estimated the proportion of mosquitoes infected at least twice to be between 2.3-3.8% in Klesso and between 0.4-0.9% in Longo.

### 4. Predicting parasite exposure in mosquitoes from prevalence in human

Gametocyte slide prevalence in human was able to predict estimates of parasite exposure with good accuracy during the rainy season in Klesso and Longo (R= 0.88, p-value= 0.033, Figure 4.a) for a relatively small dataset which had a low range of estimates of gametocyte slide positive individuals (0.01-0.08%). Malaria parasite prevalence of any parasite stage in human was also a promising predictor of parasite exposure, although only of marginal significance (R= 0.83, p-value= 0.058, figure 4. b) Gametocyte or trophozoite densities in humans were poor predictors of estimates of parasite exposure (gametocyte density: p-value= 0.658; trophozoite density: p-value= 0.419). The human-mosquito transmission probability (the probability that a mosquito becomes infected following a blood meal on a parasite carrier) was also estimated but could not be explained by the prevalence or density of either all blood stages parasites or gametocytes alone (as determined by microcopy).

**Figure 4.**
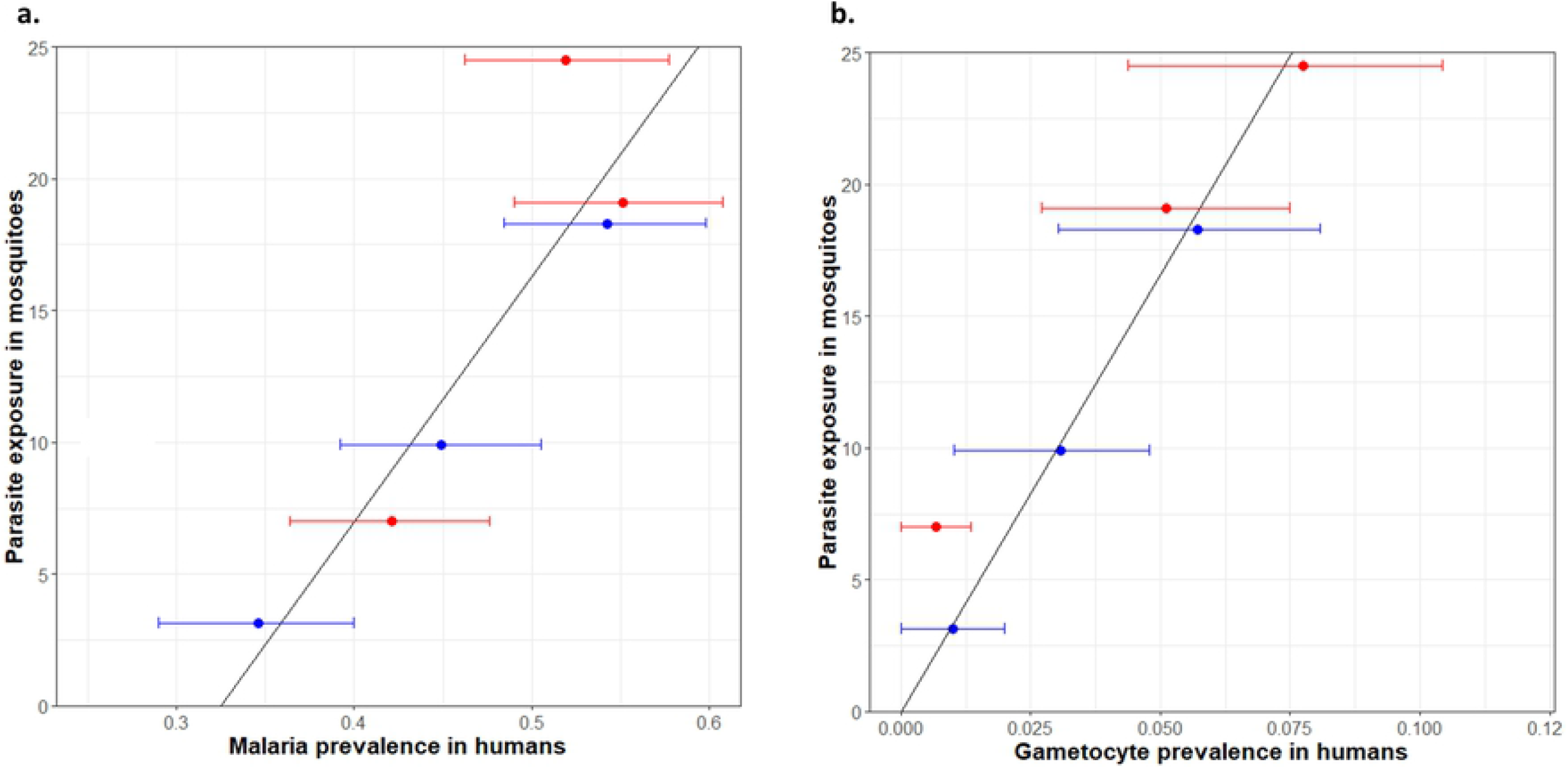
The relationship between observed prevalence of any *P. falciparum* stage (a) or gametocytes (b) estimated by microscopy in the human population and model predictions for parasite exposure (the mean number of oocysts acquired per infectious blood meal). Red points are estimates for Klesso whilst blue points are estimates for Longo for each month during the rainy season. Horizontal lines indicate 95% confidence interval estimates for malaria prevalence.

#### Cameroon

Mosquito data was collected in two villages in Cameroon (Mbelengui and Ekali) during the rainy season (May-July), and mosquitoes were maintained for 7 days in the laboratory before dissection. Prevalence and abundance of *Anopheles* mosquitoes in Cameroonian houses were significantly lower than in Burkina Faso during the peak of the rainy season (p-values<0.0001), and significantly higher in Mbelengui than in Ekali (prevalence: 41% (35-47%) vs. 22% (18-26%) p-value<0.001, abundance: 3.2% (2.6-3.8%) vs. 1.8% (1.4-2.2) mosquitoes per infested house p-value<0.001; Figure 5.a). A. *gambiae* was the most prominent species (84% (81-88%)), and contrary to Burkina Faso *Anopheles funestus* was also frequently observed in Cameroon (8% (6-11%) of all *Anopheles*, see supporting table 2). Bednet coverage was significantly lower in Cameroon than in Burkina Faso during the peak of the rainy season (p-value<0.0001; 34% (27-40%) in Ekali and 29% (25-33%) in Mbelengui) and houses with bednets had a significantly lower mosquito prevalence and abundance (prevalence −80.53% p-values <0.0001, abundance −38.04% p-value= 0.012; see supporting table 2 for further details).

**Figure 5:**
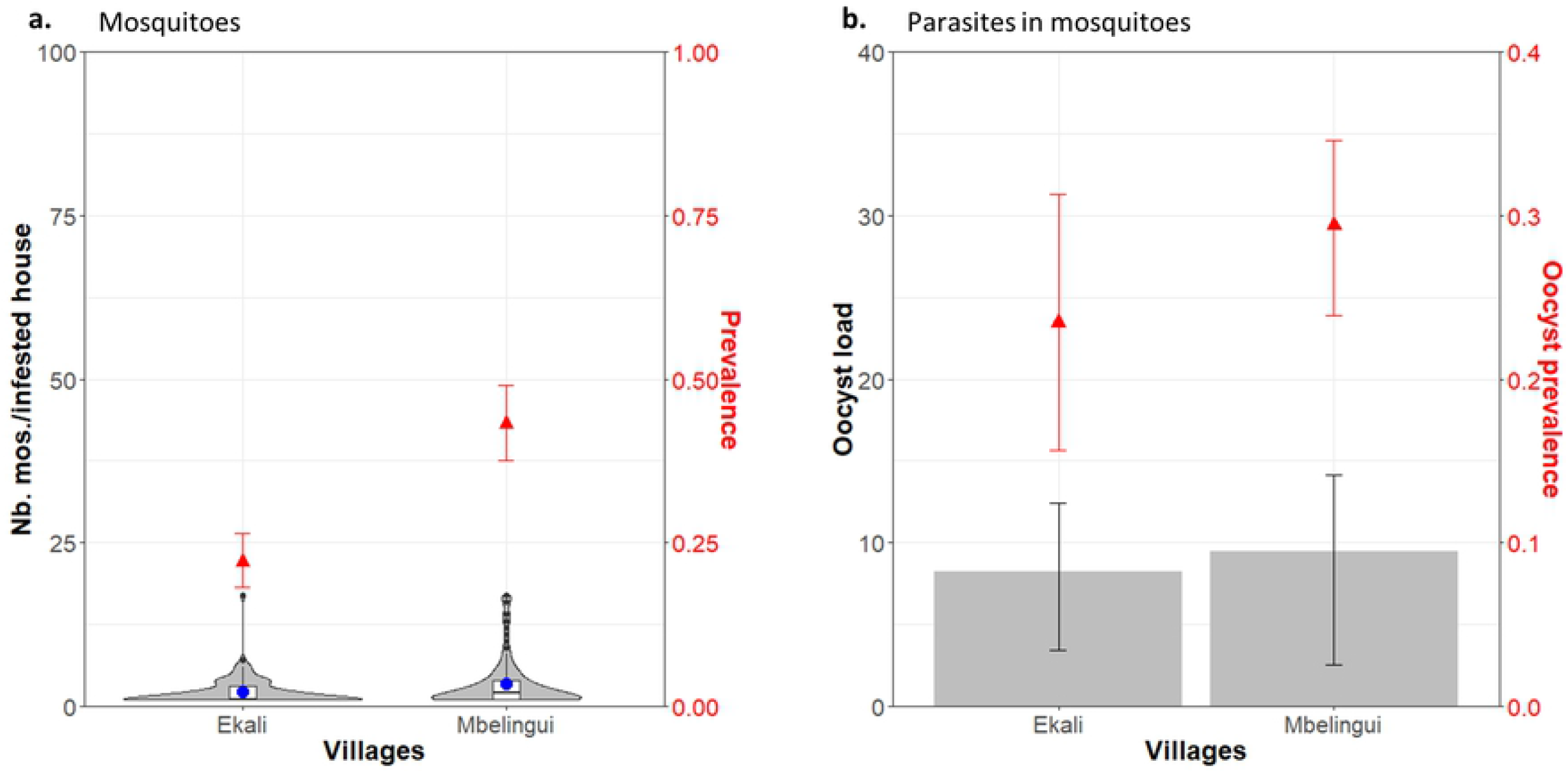
Mosquito (a) and parasite (b) data collected in Cameroon during the peak of the rainy season. a: Mosquito abundance in infested house (boxplot, left axis) and mosquito prevalence in houses (red triangles, right axis). b: Average oocyst load in infected mosquitoes dissected 7 days after collection (bars with confidence intervals) and oocyst prevalence in mosquitoes (red triangles, right axis).

Dissection showed that the prevalence of sporozoite positive mosquitoes was 42% (37-47%) 7 days after collection, higher than Burkina locations. Model estimates based on Klesso parameters (other than parasite prevalence) indicate sporozoite rates on the day of mosquito collection around 16.5% (Mbelengui) and 15% (Ekali). The prevalence oocyst positive mosquitoes was similar to Klesso during the peak of the rainy season (p-value= 0.23, Klesso: 21% (19-24%), Mbelengui: 29% (20-39%) and Ekali: 24% (7-39%)), but oocyst load in Cameroon locations was lower, with a mean 9.1% (3.2-12.9%) oocysts per infected mosquitoes (figure 5.b). Model estimates based on Klesso estimated parameters indicate Cameroon parasite exposure around 8.7 oocysts per infectious blood meal in Mbelengui and 8.1 oocysts per infectious blood meal in Ekali.

## Discussion

Wild mosquito exposure to malaria parasites in two villages in Burkina Faso was substantially higher than has previously been assumed. Overall, model estimates indicate each infectious blood meal resulted in a mean of 14 oocysts per infected mosquito. The range of parasite exposure was high, though estimates at all time points were substantially greater than was previously assumed, even in high infection areas (9,26,27). This result was driven by the high oocyst load observed in wild caught mosquitoes dissected 3 and 7 days post collection. In Burkina Faso there were on average over 10 oocysts per mosquito during the rainy season seven days after collection, which is substantially more than previous work reported (less than 5 oocysts/infected wild mosquito(24)). Estimates of sporozoite rates at day 0 (predicted using the mathematical model) were high, though were similar to another recent study from Burkina Faso(28). Observed oocyst loads were only slightly lower in Cameroon than in Burkina Faso, and consistent with a previous Cameroon study(26). This suggests that the high parasite exposure estimates in Burkina Faso are not atypical or specific to the area. There were some differences in the seasonality of parasite and mosquito abundance between the two villages in Burkina Faso (despite them being barely 100km apart) though estimates of the distribution of oocysts per mosquito and parasite exposure were broadly similar.

Gouagna’s(8) method for determining parasite exposure is to subtract the number of oocysts observed on day 3 from those observed on day 7 and therefore estimate the number of oocysts acquired the night before mosquito collection. This proved inadequate as there were significantly more oocysts at day 3 than at day 7 in some instances. Supplementary laboratory studies showed that this was due to the appearance of visible oocysts prior to day 3, with on average 60% of mosquitoes that were infected having visible oocysts on day 3. This may be due to high temperatures in the laboratories in Burkina Faso (29–31), or to a wild vector population particularly permissive to *Plasmodium* infections(32) or to other environmental factors affecting the length of the extrinsic incubation period(32). The proportion of mosquitoes with visible oocysts also appears to increase as oocyst exposure increases (supporting information 2). Temperature has previously been reported to cause both an increase in the number of oocysts per infected mosquito and a decrease in the time to oocyst appearance(31) but this is the first time (to our knowledge) that a relationship between the latter two has been observed at a constant temperature, and further studies are necessary to understand to what extent this could affect malaria parasite transmission. Biological causes for the faster appearance in more infected mosquitoes could be a higher investment in transmission stages due to oocyst competition for resources or higher mosquito immune response when infection intensity increases(32,33). This acceleration in oocyst development may result in lower numbers of sporozoites(24,34)(35) but if oocysts that become mature also break and release sporozoite earlier, mosquitoes could become infectious quicker (having a lower extrinsic incubation period) in high parasite-exposure areas, exacerbating *Plasmodium* transmission.

In Burkina Faso and in Cameroon parasite exposure was shown to be higher than is usually assumed. The percentage of mosquitoes with visible oocyst is likely to be higher in mosquitoes caught and kept in the laboratory before dissection compared to being dissected on the day of collection as mortality in the laboratory is probably lower than in the wild. Nevertheless, a previous study in Tanzania reported that only 9% of naturally infected wild mosquitoes had over 5 oocysts per mosquito when dissected on day 6 after collection(9) whereas in the present study, in Burkina, overall 65% had 1-5 oocysts, 35% had over 5 oocysts, 21% over 10 and 2% over 100 oocysts. Over-dispersion in oocyst distribution has been documented before but this is the first time a substantial proportion of mosquitoes with very high oocyst numbers has been found in nature(7,9,10,36). The occurrence of wild mosquitoes with over 100 broken oocysts indicates that it is likely these highly infected mosquitoes can survive long enough in the wild for the oocysts to release infectious sporozoites, and that even if some oocysts may fail to rupture(37,38), most do in this context. If these highly infected mosquitoes have in turn a higher sporozoite count they would be particularly infectious, as more sporozoites in salivary glands may translate into a higher probability of infection(39). However, whether sporozoite production per oocyst is maintained or decreases with oocyst load remains unclear(34,40), so it is difficult to predict whether these highly infected mosquitoes will constitute a major component of transmission and a challenge for malaria elimination.

The consequences of high parasite exposure could be far-reaching for our understanding of malaria and how transmission-reducing interventions are assessed. Results of standard and direct membrane feeding assays (SMFA and DMFA, respectively) are often challenged because the parasite exposure generated is seen as artificially high compared to natural parasite exposure(24). However, the parasite exposure we observed in high transmission area falls into a similar range as recent SMFA studies(18,41,42). Transmission-blocking interventions that show a dose-response to parasite exposure, such as transmission-blocking vaccines, might initially show lower efficacy when deployed in high-transmission areas. However, as recent results indicate, a TBV may reduce parasite exposure over time(39), increasing efficacy. This positive feedback loop could substantially increase overall effectiveness of interventions which target the parasite component of human-to-mosquito transmission. More broadly, the work has implications for the understanding of the population dynamics of *P. falciparum* and the interaction between the malaria parasite and its vector.

The experiment outlined here to determine parasite exposure is complex, requiring large numbers of mosquitoes to be kept in the laboratory and dissected. Estimates of field parasite exposure can be made by dissecting mosquitoes on the day of collection if they have not blood-fed. This does give an indication of the level of exposure, though may be biased by malaria parasite endemicity, non-human blood-feeding and any impact the parasite may have on the mosquito in the wild. Here we compare mosquito parasite exposure to the prevalence of the parasite in the human population to investigate whether this simpler to measure parameter can be used to predict exposure in the wild. Though the relationship between parasite exposure and human *Plasmodium* prevalence is particularly promising, further work is needed in more locations with differing levels of parasite endemicity. Although gametocyte prevalence is a more precise predictor of parasite exposure in our study, all-stages *Plasmodium* prevalence in humans would be an easier metric to collect and has greater ability to differentiate between parasite exposures as there is wider variability observed in the field and has and fewer sub-microscopic infections. Variation in host natural immunity between locations could also affect this relationship so more data in various epidemiological settings would be needed to corroborate the relationship. If confirmed, a human *Plasmodium* prevalence-mosquito natural parasite exposure curve could help predict initial and long-term TBV efficacy before clinical trial or deployment, as well as the minimum antibody titer necessary to achieve blockade(18). The World Health Organization has recently stressed the need to target control in high burden countries such as Burkina Faso(43). The high parasite exposure reported here may contribute to the resilience of malaria to ongoing control efforts in Burkina Faso and other high transmission settings.

## Materials & Methods

### 1. Data

Entomological and parasitological survey were performed in rural endemic areas of Burkina Faso and Cameroon, to evaluate the *Plasmodium* human-mosquito spatial-temporal transmission. Two villages in each country were served as sites for data collection: Klesso and Longo in Burkina Faso and, Mbelengui and Ekali in Cameroon. In Burkina Faso, data was collected during four different periods in 2014: the dry season (March-April), the start of the rainy season (July), the peak of the rainy season (September) and the end of the rainy season (November). In Cameroon, we were only able to collect data during one period in 2015 (May to July).

#### Mosquito collection

During each period, mosquito collection was uniformly organized: 10 people collected alive mosquitoes each morning for at least 20 days using a map highlighting the collection sites in each village. Mosquitoes were caught using a mouth aspirator in the living room of human dwellings early in the morning (44). In parallel, huts characteristics were reported, such as: presence of a bednet, type of roofing (straw, slab or metal), presence or not of paint on the walls, and localisation within locally-defined quarters. Mosquitoes were counted, *Anopheles* females were identified, pooled in a cage per village and brought to the laboratory to be maintained in the insectary. At each period, a subset of collected mosquito was dissected to identify feeding status, parturity rate and gravidity at the time of collection.

#### Mosquito rearing and dissection

Mosquito females were maintained in the insectary and dissected following Gouagna *et al.’s* work (8). For each collection date and location in Burkina Faso, mosquitoes were maintained in the laboratory (27-28°C, 70-80% relative humidity, dark-light photoperiod 12:12 hours), half of the population for 3 days and the rest for 7 days before dissection. Every mosquito midgut was dissected to count normal oocysts (as used in the results section) and report the number of broken oocysts (presented in supporting information 3). After dissection, we randomly selected 100 females from each villages and season among mosquitoes without oocyst, and 100 females among mosquitoes with oocysts. Head and thoraces of each of these mosquitoes were used for mosquito species identification via PCR (45) and sporozoite detection via qPCR (46). Sporozoite rate were then adjusted using oocyst prevalence to reflect the population rate for each season and location. In Cameroon, mosquitoes were collected and then maintained in the laboratory in similar conditions. Dissections were only carried out on day 7 only to assess sporozoite presence and oocyst number.

Parasite exposure could not be deduced from the difference in oocyst numbers between day 7 and day 3 dissections in Burkina Faso, as in Gouagna’s method(8). To understand how many parasites were acquired over a single blood meal, we conducted a supporting laboratory experiment on artificially infected mosquitoes to identify the speed at which oocysts become detectable when mosquitoes are reared in the laboratory (presented in supporting information 2), and fitted a transmission model to our data (presented section 2.2. and supporting information 3).

#### Blood collection

At the end of each mosquito collection period, blood samples were collected from volunteers within the two Burkina Faso villages. Blood smears were carried out for 300 people per time point and village (100 per age class, < 10 year old, 10-20 year old, > 20 year old), and malaria-positive people received a treatment. Gametocytes, trophozoites and schizonts were counted and the *Plasmodium* species identified by microscopy. For each blood donor, one thick and one thin smear were made on the same slide. Gametocytes and schizonts were counted against 1000 white blood cells or on up to 100 fields on the thick smear; slides were declared negative after a minimum reading of 100 fields. Trophozoites where counted against 200 white blood cells, and counting was extended to 500 white blood cells if less than 10 parasites were found. Parasite numbers were converted to parasite densities assuming 8 000 white blood cells/μl. Each slide was read three time by three independent qualified technicians, and results were compared two by two to assess concordance. If the difference between two reading results was less than half of their mean, the readings were validated and the mean of the three results was kept as parasite density. If the difference was greater between two of the three readings, the slide was re-read again until concordance.

#### Ethics statement

The Comité Ethique pour la Recherche en Santé, Burkina Faso, approved the study under ethical clearance 2014-040/MS/MRSI/CERS. Blood samples were collected from adult volunteers who provided informed consent, and from child participants with informed consent from their parent or guardian. All informed consent was written.

### 2. Analysis

The data collected was first analyzed using generalized linear mixed-effect models. Prevalence data (for mosquitoes in houses, blood stage and mosquito stage parasites) was analyzed using a binomial distribution, while intensity data (number of mosquitoes, blood stage parasite or oocysts in prevalence-positive agents) was analyzed using a zero-truncated Poisson distribution (for mosquitoes abundance in houses) or a zero-truncated negative binomial distribution (for blood and mosquito-stage parasites) with R package glmmADMB(47).

We developed a compartmental differential equation model to track mean oocyst load in the mosquito population and the proportion of susceptible, infected and infectious mosquitoes (programmed in Berkley Madonna). The model also follows oocyst prevalence and load and sporozoite prevalence in mosquitoes removed from the transmission cycle and reared for 3 to 7 days in the laboratory, to match the data collected in the field. The proportion of oocysts developed after infection is set to follow a curve depending on parasite exposure and assessed in a supplementary experiment described in supporting information 2. The model framework and equations are detailed in supporting information 3.

Berkeley Madonna optimization function was used to select the parameter values that maximized the log likelihood of the observed data (D3 and D7 sporozoite prevalence, oocyst prevalence and load) given the observed human prevalence. This was conducted for each Burkina Faso location and timepoint (see supporting information 3 for details). For Cameroon locations, parasite exposure, infection rate and D0 sporozoite rates were estimated using Klesso parameters and Cameroon sporozoite and oocyst data collected at D7.

## Acknowledgments

The work was supported by the PATH Malaria Vaccine Initiative and the UK Medical Research Council (MRC)/UK Department for International Development (DFID) under the MRC/DFID Concordat agreement. We thank the MVI team for their helpful comments on the manuscript.

